# Loss of floor plate Netrin-1 impairs midline crossing of corticospinal axons and leads to mirror movements

**DOI:** 10.1101/2020.02.20.958595

**Authors:** Oriane Pourchet, Marie-Pierre Morel, Quentin Welniarz, Nadège Sarrazin, Fabio Marti, Nicolas Heck, Cécile Galléa, Mohamed Doulazmi, Sergi Roig Puiggros, Juan Antonio Moreno-Bravo, Marie Vidailhet, Alain Trembleau, Philippe Faure, Alain Chédotal, Emmanuel Roze, Isabelle Dusart

**Author notes:** Equal contribution (Emmanuel Roze, Isabelle Dusart).

## Abstract

In human, execution of unimanual movements requires lateralized activation of the primary motor cortex, which then transmits the motor command to the contralateral hand through the crossed corticospinal tract (CST). Mutations in *NETRIN-1* alter motor control lateralization, leading to congenital mirror movements. To address the role of midline Netrin-1 on CST development and subsequent motor control, we analyzed the morphological and functional consequences of floor-plate Netrin-1 depletion in conditional knock-out mice (*Shh::cre;Ntn1*^*lox/lox*^ mice).

Here, we show that depletion of floor plate Netrin-1 critically disrupts midline crossing of the CST, whereas the other commissural systems are mostly preserved. The CST defect results in abnormal but functional ipsilateral projections, and is associated with abnormal symmetric movements. Therefore, our study reveals a new role for Netrin-1 in CST development. It also describes a unique mouse model recapitulating characteristics of human congenital mirror movements, through abnormal CST decussation.

## Introduction

Asymmetric voluntary movements, such as opening a bottle or texting a friend, are essential to everyday life. In humans, execution of unimanual movements requires communication between secondary motor areas and primary motor areas, to activate one motor cortex, which then transmits the motor command to the contralateral hand through the crossed corticospinal tract (CST). Congenital mirror movement syndrome (CMM) is a rare neurological disorder characterized by the inability to perform unilateral movements with the hands^1^. CMM patients thus experience difficulties in bimanual activities of daily living requiring asymmetric voluntary movements and fine motor coordination. Among the different defects observed in CMM patients, the most consistent one is the presence of an abnormal corticospinal tract (CST) decussation resulting in corticospinal axons projecting to both sides of the spinal cord^2^. Several mutant mice display abnormal CST decussation^3–6^. However, they do not allow the study of the contribution of abnormal CST decussation to the lateralization of motor control, because of the presence of additional guidance defects in other parts of the central nervous system (CNS). How CST development relates to the lateralization of motor control thus remains unclear.

*NETRIN-1* (*NTN1*) has been associated to CMM^7^. This gene encodes an extracellular protein, which influences the formation and the maintenance of multiple tissues^8^. In the developing nervous system, Netrin-1 acts as a guidance cue required for midline crossing of many commissural axons^9^. *Ntn1* has also been proposed to be involved in midline crossing of the CST in mice^5^. However, this hypothesis has not been confirmed because *Ntn1^KO^* mice die at birth^9^, when corticospinal axons reach the pyramidal decussation^10^. *Shh::cre;Ntn1*^*lox/lox*^ mice is a viable mouse model, in which *Ntn1* depletion is restricted to the floor plate in the regions of the CNS located caudal to the midbrain/hindbrain boundary. Interestingly, midline crossing of spinal cord and hindbrain commissural axons is mostly preserved in this model^11–14^. Yet, development of the corticospinal axons, which occurs over a protracted period, has not been investigated in these mice.

In the present study, we analyzed the anatomy of the corticospinal tract and the presence of symmetric movement in *Shh::cre;Ntn1*^*lox/lox*^ mice to determine the contribution of floor plate Netrin-1 in midline crossing of corticospinal axons and lateralization of motor control.

## Results

### Guidance of the CST is disrupted at the pyramidal decussation in *Shh::cre;Ntn1*^*lox/lox*^ mice

We first used PKCγ immunochemistry to characterize the anatomy of the CST in adult *Shh::cre;Ntn1*^*lox/lox*^ mice (Fig. 1a). The tract was comparable to control mice (Fig. 1a, Supplementary Fig. 1a) down to the medulla level, where corticospinal axons formed two compact ventromedial tracts. In control mice, the CST remained fasciculated until the junction between the brainstem and the spinal cord. There, it turned dorsally and crossed the midline to enter into the dorsal funiculi of the spinal cord, forming the so-called pyramidal decussation. In *Shh::cre;Ntn1*^*lox/lox*^ mice, the corticospinal axons defasciculated prematurely just before the junction between brainstem and spinal cord. At this level, the CST spread laterally and the pyramidal decussation was thinner than in the control. Accordingly, fewer corticospinal axons were detected in the dorsal funiculi of the spinal cord of *Shh::cre;Ntn1*^*lox/lox*^ mice and the remaining corticospinal axons were observed in the ventromedial and lateral funiculi of the spinal cord. To assess the laterality of these abnormal corticospinal projections, we injected either tdTomato-expressing adenovirus or biotinylated dextran amine unilaterally in the motor cortex of *Shh::cre;Ntn1*^*lox/lox*^ and control mice (Fig. 1b, Supplementary Fig. 1b). All labelled corticospinal axons reached the contralateral side of spinal cord in control mice (Fig. 1c, Supplementary movie 1 and Supplementary Fig. 1c). In *Shh::cre;Ntn1*^*lox/lox*^ mice, corticospinal axons were detected on both sides of the spinal cord with a variable proportion of axons on each side (Fig. 1c, Supplementary movie 2 and 3). FP-Netrin-1 is thus required for midline crossing of corticospinal axons at the pyramidal decussation.

**Figure 1:**
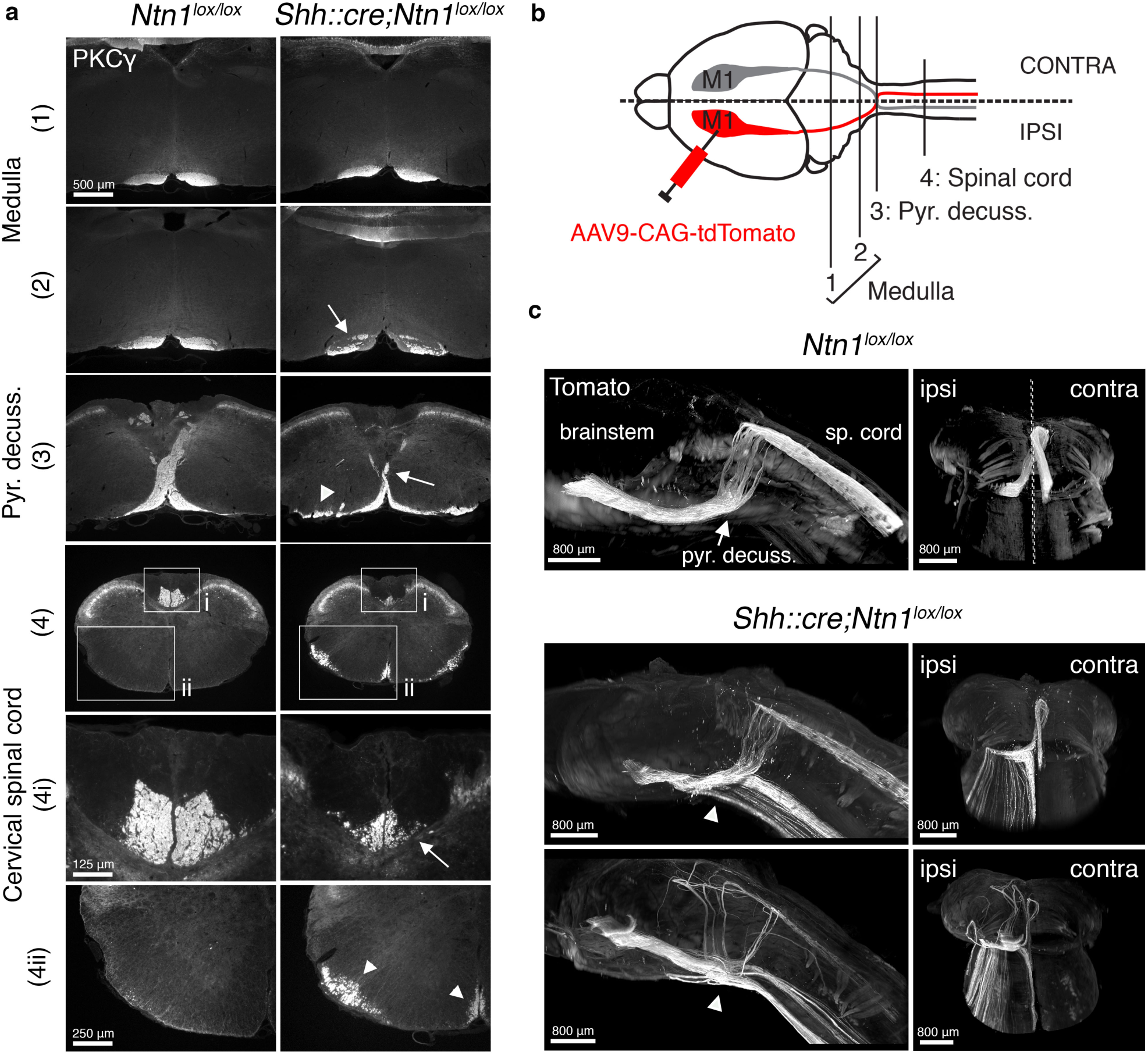
Guidance of the corticospinal tract (CST) is disrupted at the pyramidal decussation in *Shh::cre;Ntn1*^*lox/lox*^ mice. **a**, Coronal sections of adult *Ntn1I*^*lox/lox*^ (n = 4) and *Shh::cre;Ntn1I*^*ox/lox*^ mice (n=5) stained with anti-PKCγ, a CST marker, at different levels: (1) upper and (2) lower medulla, (3) pyramidal decussation, and (4) cervical spinal cord. The CST of *Shh::cre;Ntn1I*^*ox/lox*^ mice is similar to control in (1), but less fasciculated (arrow) in (2). In (3), pyramidal decussation is thinner (arrow) and the CST spreads laterally (arrowhead). In (4), less axons are detected in the dorsal funiculi (4i, arrow) and some axons are seen at ectopic positions in the ventromedial and lateral funiculi (4ii, arrowheads). **b**, Schematic showing the CNS levels (1, 2, 3, 4) for the coronal sections presented in a and CST tracing strategy. **c**, Three-quarter (left pannel) and caudal (right pannel) 3D views of cleared brainstems of a *Ntn1I*^*ox/lox*^ adult mouse (top) and two *Shh::cre;Ntn1I*^*ox/lox*^ mice (bottom). The CST from the left motor cortex is labelled with tdTomato. In *Ntn1I*^*lox/lox*^ *mice* (n = 2), at the pyramidal decussation (arrow), all labeled corticospinal axons turn dorsally and cross the midline (dotted line) to project into the contralateral dorsal funiculus. In *Shh::cre;Ntn1*^*lox/lox*^ mice (n=3), many corticospinal axons remain ventral (arrowheads), do not cross the midline, and project aberrantly in the ipsilateral ventral spinal cord. Some axons turn dorsally without crossing the midline and project aberrantly into the ipsilateral dorsal funiculus (bottom pannel). pyr. decuss.= pyramidal decussation; sp. cord= spinal cord; ipsi, contra are respectively ipsilateral and contralateral sides with respect to the injection side.

### Lateralization of corticospinal projections is altered in the spinal cord of *Shh::cre;Ntn1*^*lox/lox*^ mice

We then analyzed how such disruption of CST guidance at the pyramidal decussation affects the pathfinding and connectivity of CST axons (Fig. 2a). In the mouse spinal cord, corticospinal axons leave the white matter to enter the grey matter, mostly at the cervical and lumbar enlargements^10^. There, they terminate mainly in the dorsal and intermediate horns of the contralateral spinal cord, where they contact interneurons, which eventually transmit the command to forelimb and hindlimb motoneurons^15,16^. At both cervical and lumbar enlargements, the total number of corticospinal axons detected in the white matter was similar between *Shh::cre;Ntn1*^*lox/lox*^ and control mice (Fig. 2a,b). However, most of the corticospinal axons were detected in the ipsilateral side of the spinal cord in *Shh::cre;Ntn1*^*lox/lox*^ mice, whereas this proportion did not exceed 5% in control mice. To assess the density of corticospinal synapses in the grey matter, we studied the colocalization of VGLUT1 puncta with tomato positive boutons (Fig. 2c). In control mice, corticospinal synapses were almost exclusively found in the contralateral grey matter, with a higher density in the dorsal horn compared to the intermediate horn. In *Shh::cre;Ntn1*^*lox/lox*^ mice, synapses were detected on both sides of the grey matter, with a proportion of synapses in the ipsilateral side ranging from 24 to 70%. Yet, the dorso-ventral distribution of synapses was preserved in these mice.

**Figure 2:**
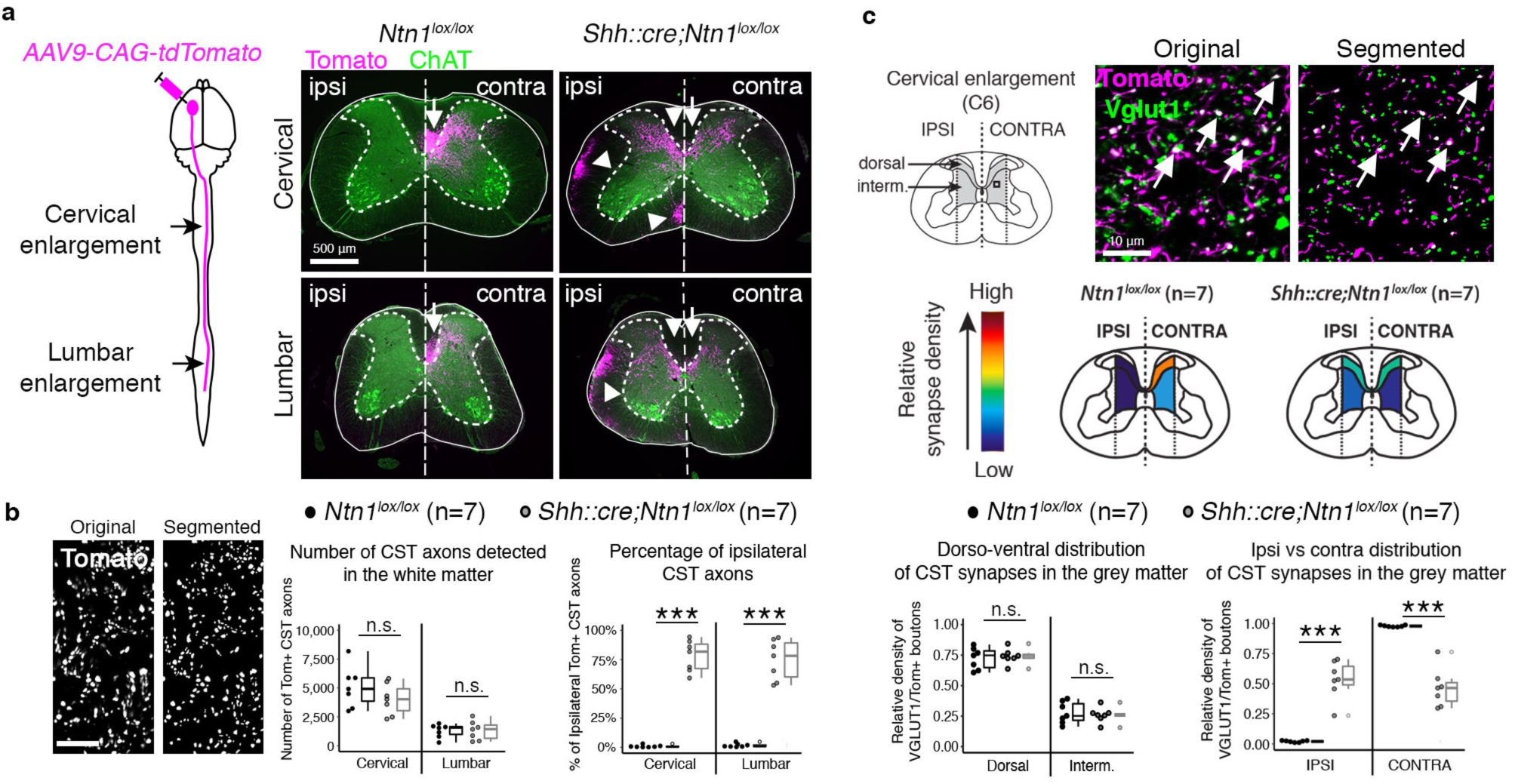
Anatomical caracteristics of the aberrant corticospinal projections in *Shh::cre;Ntn1*^*lox/lox*^ mice. **a**, Schematic representing unilateral CST labelling and coronal sections of adult *Ntn1*^*lox/lox*^ (n=7) and *Shh::cre;Ntn1*^*lox/lox*^ mice (n=7) stained with anti-DsRED (revealing Tomato) and anti-ChAT (Choline AcetylTransferase, motoneuron marker). Corticospinal axons are detected in the contralateral dorsal funiculus (arrow) in *Ntn1*^*lox/lox*^ mice, and in contralateral dorsal (arrow) and ipsilateral dorsal, ventral and lateral funiculi (arrowheads) in *Shh::cre;Ntn1*^*lox/lox*^ mice. In grey matter, corticospinal axons innervate mostly contralateral dorsal and intermediate regions in *Ntn1*^*lox/lox*^ and both contralateral and ipsilateral regions in *Shh::cre;Ntn1*^*lox/lox*^ mouse. **b**, Original and segmented confocal images of a subregion of a *Shh::cre;Ntn1*^*lox/lox*^ dorsal funiculus: axons are detected individually; Number of labelled corticospinal axons in the white matter, P_Cervical_=0.231, P_Lombar_=0.938 (two-sided-t-test); Proportion of ipsilateral axons, ***P_Cervical_=0.001 (Mann Whitney), ***P_Lombar_<0.0001 (two-sided-t-test). **c**, Schematic of a cervical spinal cord coronal section showing in grey the region of interests (ROIs) where the density of corticospinal synapses was analysed; Original and segmented confocal images of a subregion of *Shh::cre;Ntn1*^*lox/lox*^ grey matter: corticospinal synapses (white points, arrows) are identified by the colocalization of anti-DsRED (revealing Tomato) and anti-VGLUT1; Color-coded diagram representing the average distribution of the corticospinal synapse density among the 4 ROls of the grey matter; Relative density of corticospinal synapse in dorsal and intermediate regions, P=0.633 (two-sided-t-test). Relative density in ipsilateral and contralateral sides, ***P<0.0001 (two-sided-t-test). Original: original image, Segmented: image obtained after automated segmentation; Interm.: Intermediate; Data are presented as scatter and box plot, with first and third quartiles, and median.

### The aberrant ispilateral corticospinal projections are functional in *Shh::cre;Ntn1*^*lox/lox*^ mice

Next, we analyzed the functional consequences of the abnormal lateralization of the CST on cortico-motoneuronal pathway activity. We applied unilateral cortical electric stimulations and recorded the evoked electromyographic (EMG) responses in contralateral and ispilateral gastrocnemii muscles (Fig. 3a). In control mice, cortical stimulation triggered contraction of the contralateral gastrocnemius with corresponding EMG responses (Fig. 3a-c). In *Shh::cre;Ntn1*^*lox/lox*^ mice, similar cortical stimulations elicited either contralateral, bilateral or ipsilateral EMG responses, in variable proportions (Fig. 3a-c). This shows that aberrant ipsilateral corticospinal projections of *Shh::cre;Ntn1*^*lox/lox*^ mice are functional. The ipsilateral only EMG responses produced by cortical stimulations (Fig 3a, bottom right) further suggests that the ipsilateral projections are not collateral branches of the contralateral corticospinal neurons.

**Figure 3:**
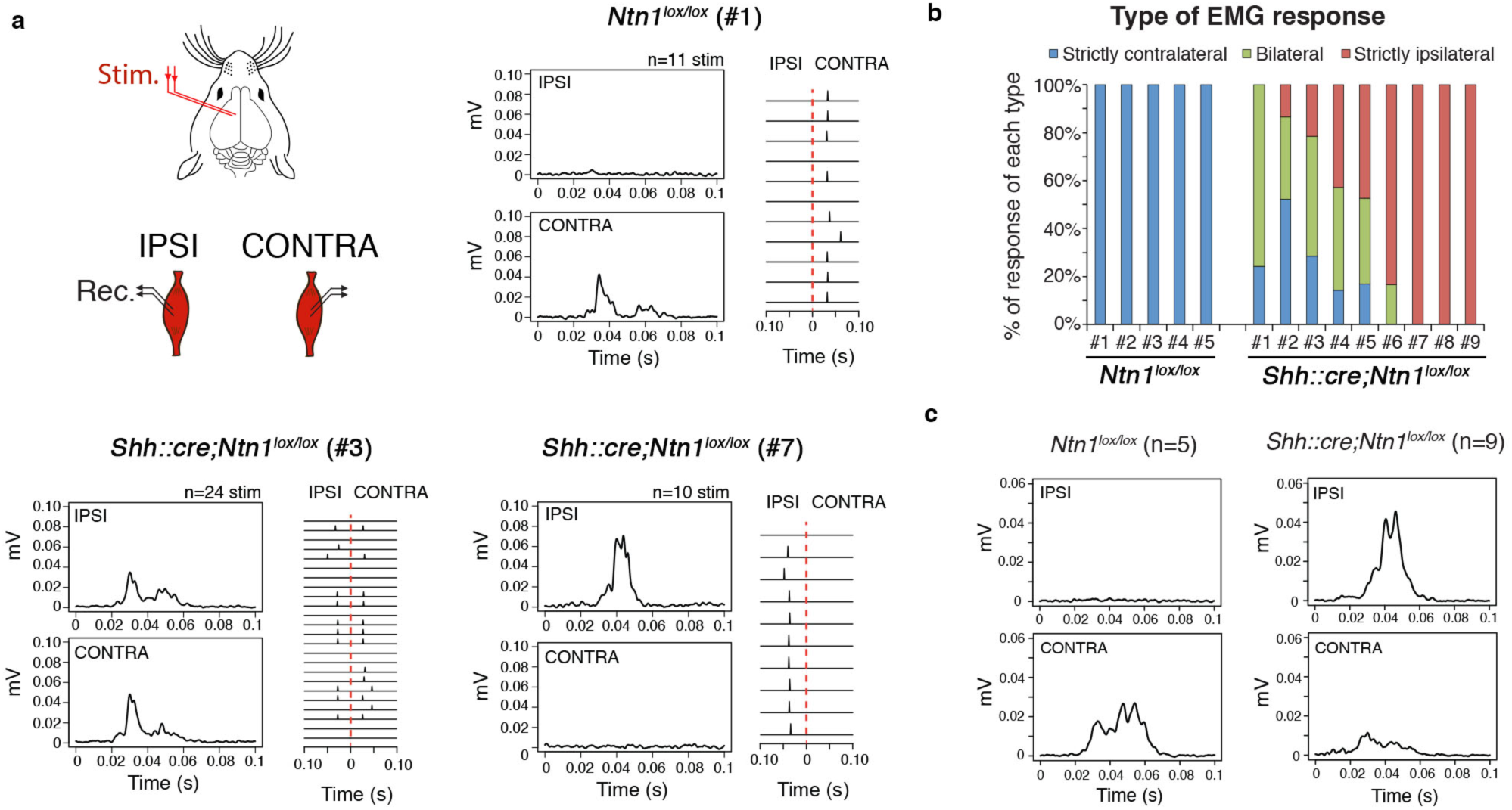
Functional characteristics of the aberrant corticospinal projections in *Shh::cre;Ntn1*^*lox/lox*^ mice. **a**, Schematic of the electrophysiological procedure. Electric stimulations were delivered in one motor cortex and evoked EMG were recorded in ipsilateral and contralateral gastrocnemii muscles of adult *Ntn1*^*lox/lox*^ (n=5) and *Shh::cre;Ntn1*^*lox/lox*^ mice (n=9). Examples of average EMG responses from 3 mice (#1 *Ntn1*^*lox/lox*^, #3 and #7 *Shh::cre;Ntn1*^*lox/lox*^ and corresponding raster plots showing different patterns of reponses. **b**, Unilateral cortical stimulations evoked ipsilateral, bilateral and/or contralateral EMG responses in *Shh::cre;Ntn1*^*lox/lox*^ mice and only contralateral EMG responses in *Ntn1*^*lox/lox*^ mice. **c**, Average EMG responses for the 5 *Ntn1*^*lox/lox*^ (left) and the 9 *Shh::cre;Ntn1*^*lox/lox*^ mice (right). Stim.= Stimulation; Rec.=Recording.

### Abnormalities of motor tracts are restricted to the CST in *Shh::cre;Ntn1*^*lox/lox*^ mice

In *Shh::cre;Ntn1*^*lox/lox*^ mice, disruption of the CST decussation results in abnormal but functional ipsilateral projections. Previous characterization of *Shh::cre;Ntn1*^*lox/lox*^ mice revealed that guidance of commissural axons in the spinal cord was mostly preserved^11–14^. In *Shh::cre;Ntn1*^*lox/lox*^ mice, we found no alteration of *Ntn1* expression in the forebrain (Supplementary Fig. 2). Using unilateral injection of a retrograde tracer in the cervical spinal cord (Fig. 4a,b), we analyzed rubrospinal and reticulospinal tracts in *Shh::cre;Ntn1*^*lox/lox*^ mice, two descending tracts known to contribute to motor control^17^. While we confirmed the lateralization abnormalities of the CST, we found no significant differences in the lateralization of the rubrospinal and reticulospinal tracts between *Shh::cre;Ntn1*^*lox/lox*^ and control mice (Fig. 4c-e). These mice are therefore a unique model to analyze the contribution of CST decussation to the lateralization of motor control.

**Figure 4:**
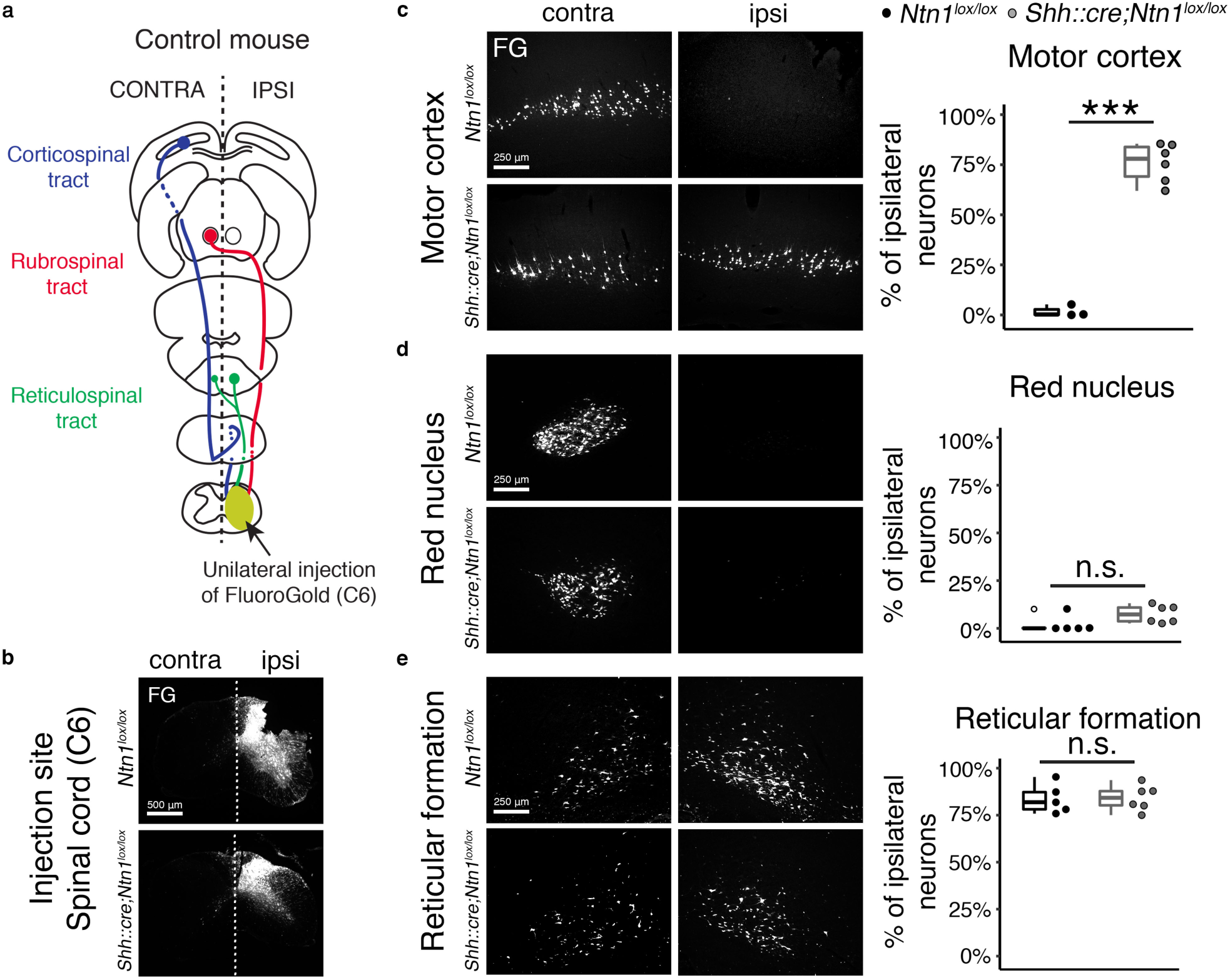
Lateralization of rubrospinal and reticulospinal tracts is preserved in *Shh::cre;Ntn1*^*lox/lox*^ mice. **a**, Schematic showing the injection site of the retrograde tracer (yellow ellipse) and the projection and origin of the corticospinal (blue), rubrospinal (red) and reticulospinal tracts (green) in an adult control mouse. **b**, Coronal sections showing the unilateral injection of FluoroGold on the right side of the lower cervical spinal cord (C6) in *Ntn1*^*lox/lox*^ (n=5) and *Shh::cre;Ntn1*^*lox/lox*^ mice (n=6). **c**, Left: Coronal sections of the contralateral and ipsilateral motor cortices of *Ntn1*^*lox/lox*^ (n=3) and *Shh::cre;Ntn1*^*lox/lox*^ mice (n=6). FluoroGold-positive neurons can only be detected on the contralateral side in *Ntn1*^*lox/lox*^ mice whereas they are detected on both sides in *Shh::cre;Ntn1*^*lox/lox*^ mice. Right: *Ntn1*^*lox/lox*^ (black dots) and *Shh::cre;Ntn1*^*lox/lox*^ mice (grey dots). ***P<0.001 (two-sided t test). **d**, Coronal sections of the contralateral and ipsilateral red nuclei of *Ntn1*^*lox/lox*^ (n=5) and *Shh::cre;Ntn1*^*lox/lox*^ mice (n=6). Almost all FluoroGold-positive neurons are detected on the contralateral side in both *Ntn1*^*lox/lox*^ and *Shh::cre;Ntn1*^*lox/lox*^ mice. P=0.1173 (Mann-Whitney). **e**, Coronal sections of the contralateral and ipsilateral medullary reticular formation of *Ntn1*^*lox/lox*^ (n=5) and *Shh::cre;Ntn1*^*lox/lox*^ mice (n=6). FluoroGold-positive neurons are detected on both sides, with a higher proportion of labeled neurons on the ipsilateral side for both *Ntn1*^*lox/lox*^ and *Shh::cre;Ntn1*^*lox/lox*^ mice. P=0.9148 (two-sided t test). FG: FluoroGold; ipsi, contra are respectively ipsilateral and contralateral sides with respect to the injection side. Data are presented as scatter and box plot, with first and third quartiles, and median.

### Lateralization of voluntary movements is altered in *Shh::cre;Ntn1*^*lox/lox*^ mice

We first submitted *Shh::cre;Ntn1*^*lox/lox*^ mice to classic motor tests. Their general motor behavior was mostly undistinguishable from control littermates (Fig. 5a), indicating that changes in ratio of contralateral vs. ipsilateral corticospinal projections does not significantly influence basic overground locomotion, balance or motor coordination. We then analyzed the ability of *Shh::cre;Ntn1*^*lox/lox*^ mice to perform lateralized coordinated movements in two specific contexts, namely automatic movements and voluntary movements. Lateralization during automatic movements was investigated with the treadmill locomotion task. In this task, left-right alternation of forelimbs and hindlimbs was preserved in *Shh::cre;Ntn1*^*lox/lox*^ mice. The proportion of symmetric movements was very low and similar to control mice (Fig. 5b, Supplementary movies 4,5). We then used the adaptive locomotion and the exploratory reaching tasks to analyze lateralized motor control during voluntary movements. These two tests have been developed specifically to reveal the presence of abnormal symmetric voluntary movements in mice^18^. Interestingly, *Shh::cre;Ntn1*^*lox/lox*^ mice produced a significantly higher proportion of symmetric movements compared to control mice, both in adaptive locomotion and reaching tasks (Fig. 5c,d and Supplementary movies 6-9). Overall, these findings reveal that lateralized control of voluntary movements is selectively impaired in *Shh::cre;Ntn1*^*lox/lox*^ mice.

**Figure 5:**
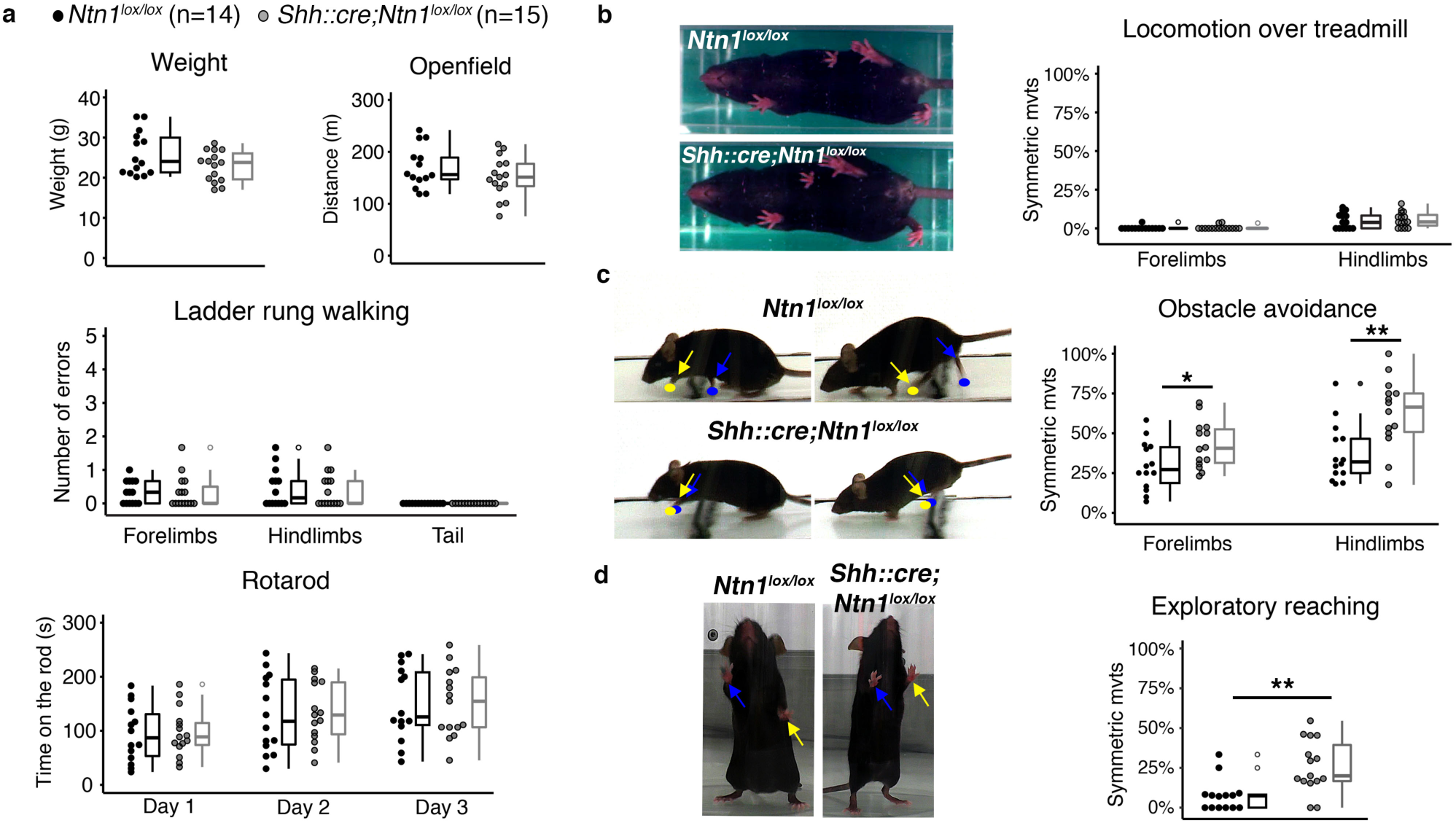
Lateralization of voluntary movements is altered in Shh::cre;Ntn1 ^*lox/lox*^ mice. **a**, *Ntn1*^*lox/lox*^ (black dots, n= 14) and *Shh::cre;Ntn1*^*lox/lox*^ mice (grey dots, n= 15) have similar weight. P= 0.084 (two-sided-t-test). The general motor behavior of *Shh::cre;Ntn1*^*lox/lox*^ is not impaired. They perform as well as their control littermates in the openfield, on the ladder rung task and on the rotarod. P_Openfield_= 0.228 (two-sided-t-test), P_Ladder-Forelimbs_= 0.7151 (Mann-Whitney), P_Ladder-Hindlimbs_= 0.8471 (Mann-Whitney), P_Ladder-Tail_= 1 (Mann-Whitney), P_Rotarod-Day1_= 0.911; P_Rotarod-Day2_= 0.874; P_Rotarod-Day3_= 0.966 (two-sided-t-test). **b**, Left: Picture of a Ntn1lox/lox mouse (top) and a *Shh::cre;Ntn1*^*lox/lox*^ mouse (bottom) performing alternate movements with both forelimbs and hindlimbs during treadmill locomotion. Right: Low proportion of symmetric strides in *Shh::cre;Ntn1*^*lox/lox*^ and *Ntn1*^*lox/lox*^ mice. P_Forelimbs_= 0.8131, P_Hindlimbs_= 0.7151 (Mann-Whitney). **c**, Left top: Picture of a *Ntn1*^*lox/lox*^ mouse avoiding the obstacle with alternate movements of the forelimbs and hindlimbs. Left bottom: Picture of a *Shh::cre;Ntn1*^lox/lox^ mouse avoiding the obstacle with synchronous movements of the forelimbs and hindlimbs. Yellow and blue dots/arrows indicate the positions of left and right fore- and hindpaws respectively. Right: In the obstacle avoidance test, *Shh::cre;Ntn1*^*lox/lox*^ mice perfom more symmetric movements with the forelimbs and the hindlimbs. *P_Forelimbs_=0.03, **P_Hindlimbs_= 0.003 (two-sided t test). **d**, Left: Pictures of a *Ntn1*^*lox/lox*^ mouse and a *Shh::cre;Ntn1*^*lox/lox*^ mouse performing respectively alternate or synchronous exploratory movements with the forepaws. Yellow and blue arrows indicate the positions of the left and right forepaws respectively. Right: In the exploratory reaching task, *Shh::cre;Ntn1*^*lox/lox*^ mice perform more symmetric movements than controls. **P=0.0021 (Mann-Whitney). Data are presented as scatter and box plot, with first and third quartiles, and median.

## Discussion

Using the viable and recently generated *Shh::cre;Ntn1*^*lox/lox*^ mouse model, we demonstrated that midline crossing of corticospinal axons is disrupted in the absence of FP-Netrin-1. This unveils a distinct and unique guidance mechanism for corticospinal axons, since FP-Netrin-1 was proven to be dispensable or mostly dispensable for midline crossing of hindbrain commissural axons and spinal commissural axons respectively. Guidance of these ventral commissural axons, which occurs during early embryonic stages, relies mainly on another source of Netrin-1, produced by the progenitor cells of the ventricular zone^11–14^. Interestingly, at the time when corticospinal axons cross the CNS midline (around birth), these progenitor cells are virtually absent^19^. FP-Netrin-1 may therefore constitute the main source of Netrin-1 for corticospinal axons in the caudal hindbrain. This could explain its major involvement in the guidance of CST axons.

Several mechanisms could account for the guidance defect of CST axons in *Shh::cre;Ntn1*^*lox/lox*^ mice. FP-Netrin-1 could act directly on corticospinal axons through one of the Netrin-1 receptors. Interestingly, mutations in the Netrin-1 receptors Dcc or Unc5c also disrupt midline crossing of corticospinal axons^5^. However, expression of these genes in the cortical layer V is very low at the time when the CST crosses the midline^5,20^ and specific depletion of Dcc in the CST does not impair pyramidal decussation^21^. A direct effect of FP-Netrin-1 would therefore involve Netrin-1 receptors other than Dcc or Unc5c. Alternatively, depletion of Netrin-1 from the FP, or from the most ventral part of the ventricular zone in the rostral midbrain (see Supplementary Fig. 3) could indirectly influence the pyramidal decussation, by inducing subtle morphological or molecular alterations along the CST trajectory.

We found that disrupting the pyramidal decussation does not abrogate the further development of misguided corticospinal axons, which follow alternative routes in the spinal cord. These alternate trajectories might correspond to secondary or transitory routes for corticospinal branches during development^22,23^. Strikingly, the innervation pattern of the grey matter was relatively well preserved despite the major alteration of CST lateralization and trajectories within the spinal cord. In line with these observations, our electrophysiological recordings further demonstrated that the misguided uncrossed corticospinal projections were functional.

We show that a specific alteration of CST decussation in a mouse model is sufficient to recapitulate CMM phenotype, i.e. abnormal symmetric movements in voluntary tasks but not in locomotion. The *Shh::cre;Ntn1*^*lox/lox*^ mice did not display any “hopping” behavior, i.e. synchronized activity of the homologous limbs during locomotion. The absence of hopping in *Shh::cre;Ntn1*^*lox/lox*^ mice during treadmill locomotion demonstrates that corticospinal projections are not involved in the lateralization of motor control during automatic behaviors, in keeping with previous findings^18^. The hopping behavior, reported in *Dcc*^*Kanga*^ mice carrying a spontaneous viable mutation in Dcc^5^, is indeed caused by an impairment of the spinal circuitry^24^.

In humans, the three main disorders associated with abnormal pyramidal decussation are CMM, X-linked Kallmann syndrome and Klippel–Feil syndrome^16^. Mirror movements are the common clinical sign of these syndromes and represent the only clinical manifestation of CMM. The anatomical and functional characterization of the corticospinal projections in our mouse model further indicates that the aberrant CST from one motor cortex projects either to the contralateral or to the ipsilateral side of the spinal cord rather than bilaterally, and might not *per se* be able to transmit the motor command in a strictly symmetric manner. Interestingly, CMM patients have an abnormal bilateral pattern of activation in cortical motor areas during unilateral movement execution and preparation^25,26^. We hypothesize that this abnormal activation pattern could reflect a strategy for adapting to aberrant CST projections, eventually resulting in the generation of mirror movements.

In summary, our work reveals a unique guidance mechanism for corticospinal axon midline crossing, with a critical dependence on floor plate Netrin-1. Disruption at the pyramidal decussation does not prevent further development of misguided corticospinal axons, which form an abnormal but functional ipsilateral tract. These aberrant corticospinal projections are associated to a selective increase of symmetric movements that can be related to mirror movements described in human CMM. To date, *Shh::cre;Ntn1*^*lox/lox*^ mice are the only mouse model that recapitulates both the anatomical and behavioral impairment of CMM.

## Materials and Methods

### Mouse lines and genotyping

*Shh::cre* (Jackson laboratory) and *Ntn1*^*lox*^ (given by P. Melhen, Lyon) were previously described in ^27^ and ^11^ respectively. They were maintained on a C57BL/6J background and were genotyped by PCR as described previously. All animal procedures were approved by the Regional Animal Experimentation Ethics Committee C2EA-05 Charles Darwin and the French Ministère de l’éducation nationale de l’enseignement supérieur et de la recherche (projects N°01558.03, 2018042815492966). We strictly performed these approved procedures.

### Surgeries and tracer injections

Buprenorphine (Buprecare) was injected subcutaneously 30 min before the surgery at 0.1mg/kg. During surgeries and injections, mice were anesthetized with isoflurane and body temperature was maintained at 37°C by a heating pad.

#### Anterograde tracing of the CST with AAV9-CAG-tdTomato

2-month old adult mice (7 *Ntn1*^*lox/lox*^ and 7 *Shh::cre;Ntn1*^*lox/lox*^ mice) were placed in a stereotaxic frame (Kopf Instrument) and lidocaïne (Xylovet) was injected subcutaneously at 3mg/kg 5 min before the surgery. AAV9-CAG-tdTomato (MIRCen) was diluted in saline to a final titer of 1.7.10^13^ GC/mL. Four injections of 0.5 μL of this AAV solution were administered at 1.6 nL/s in the left motor cortex using the following coordinates: 0.75mm rostral to Bregma, 1 and 2 mm lateral, at a depth of 1 mm for the 2 first injections and 0.25 mm caudal to Bregma, 1 and 2 mm lateral, at a depth of 1 mm for the 2 last injections. At each injection point, the needle was left in place for 2 min before and 4 min after the injection to avoid leakage. After surgery, the wound was cleansed and the skin sutured. Analysis was carried out 3 weeks after injection.

#### Anterograde tracing of the CST with BDA

Unilateral injections of biotinylated dextran amine (BDA, MW 10 000, SIGMA) were performed in the left motor cortex of 2-month old adult mice (4 *Ntn1*^*lox/lox*^, 1 *Shh::cre*, 1 *Shh::cre;Ntn1*^*lox/*+^ and 5 *Shh::cre;Ntn1*^*lox/lox*^ mice) following a procedure similar to the AAV9-CAG-tdTomato injection procedure. BDA was diluted in saline to a final concentration of 10%. Six injections of 0.2 μL were administered at 1.6 nL/s in the left motor cortex using the following coordinates: 1mm rostral to Bregma, 1 and 2 mm lateral, at a depth of 1 mm for the first couple of injections, 0.25mm caudal to Bregma, 1 and 2 mm lateral, at a depth of 1 mm for the 2 second couple of injections and 1mm caudal to Bregma, 1 and 2 mm lateral, at a depth of 1 mm for the last couple of injections. At each injection point, the needle was left in place for 3 min before and after the injection to avoid leakage. After surgery, the wound was cleansed and the skin sutured. Analysis was carried out 2 weeks after injection.

#### Retrograde tracing of descending tracts with Fluorogold

Retrograde tracing was performed on 2-month-old adult mice (13 *Ntn1*^*lox/lox*^ and 18 *Shh::cre;Ntn1*^*lox/lox*^ mice in total). The lower cervical spinal cord was exposed and 0.4-0.6 μL of a 4% Fluorogold (Fluorochrome) solution in 0.9% saline was injected at 2 nL/s on the right side of C6-C7. After the injection, the needle was kept in place for an additional 3 min to avoid leakage of tracer. The wound was closed by suturing the paraspinal muscles and skin. Carprofène (rimadyl) was injected subcutaneously at 5mg/kg on the wound and Buprenorphine (Buprecare, 1mg/kg per os) was added to the drinking water. Analysis was carried out 2 weeks after injection.

In all cases, mice were killed by an intraperitoneal injection of Euthasol (150mg/kg) followed by perfusion with 4% paraformaldehyde in 0.12 M phosphate buffer, pH 7.4 (PFA).

### 2D analysis of brain and spinal cord

After perfusion, brains and spinal cords were collected and post-fixed in the same fixative for 4 h at 4 °C. Samples were cryoprotected in a solution of 20% sucrose in 0.12 M phosphate buffer (pH 7.4). Brains were frozen in isopentane at −35°C, and stored at −80°C until processing. Spinal cords were embedded in Tissue-Teck OCT (VWR, Fontenay sous bois, France), frozen in isopentane at −30 °C and stored at −20 °C until processing. Samples were cut at 30 μm in coronal plane with a cryostat (Leica Microsystems) and serially mounted on slides.

### Immunohistochemistry and revelation of BDA

Sections were washed in PBS and blocked for 1 h with a PBSGT solution (1X PBS, 0.2% gelatin, 0.25% Triton X-100) containing 0.1 M lysine (Merck). Sections were then incubated overnight at room temperature in the PBSGT solution with the following primary antibody: rabbit anti-PKCγ (1/100, Santa Cruz, sc-211), guinea-pig anti-VGLUT1 (1/3000, Millipore, AB5905), goat anti-Chat (1/300, Millipore, AB144P), rabbit anti-DsRED (1/500, Living Colors, 632496), followed by 2h incubation in species-specific secondary antibody directly conjugated to fluorophores (Cy3 or Cy5 from Jackson ImmunoResearch or Alexa Fluor 488 from Invitrogen). Sections were mounted in Mowiol (Calbiochem, La Jolla, CA). BDA labelling was revealed by streptavidin conjugated to the Cy5 fluorophore (1/400, Invitrogen) that was added to the primary antibody solution.

### In situ hybridization (ISH)

#### Riboprobe synthesis

cDNAs encoding mouse *Ntn1* exon 3 were used. The *in vitro* transcription was carried out by using the Promega kit (Promega, Charbonnières, France) and probes were labeled with digoxigenin-11-d-UTP (Roche Diagnostics, Indianapolis, IN). *Tissue preparation:* E13 embryos (3 *Ntn1*^*lox/lox*^ and 3 *Shh::cre;Ntn1*^*lox/lox*^) and E14 embryos (3 *Ntn1*^*lox/lox*^ and 3 *Shh::cre;Ntn1*^*lox/lox*^) were fixed by immersion in 4% PFA over night at 4°C. Samples were cryoprotected in a solution of 20% sucrose in 0.12M phosphate buffer (pH7.4), frozen in isopentane at −35°C and then cut in coronal plane at 20μm with a cryostat (Leica).

#### In situ hybridization protocol

Brain sections were postfixed for 10 min in 4% PFA, washed in PBS, treated with proteinase K (20μg/ml; Invitrogen) for 2 min, postfixed for 5 min in 4% PFA, washed in PBS, acetylated, washed in PBS 1% Triton X-100. Slides were incubated 2 hours at room temperature with hybridization buffer (50% formamide, 5x SSC, 5x Denhardt’s, 250 μg/ml yeast tRNA, and 500 μg/ml herring sperm DNA, pH7.4). Then, tissue sections were hybridized overnight at 72°C with riboprobes (1/200). After hybridization, sections were rinsed for 2 hours in 0.2x SSC at 72°C, and blocked in 0.1 M Tris, pH 7.5, 0.15 M NaCl (B1) containing 10% normal goat serum (NGS) for 1 hour at room temperature. After blocking, slides were incubated overnight at room temperature with anti-DIG antibody conjugated with the alkaline phosphatase (1/5,000, Roche Diagnostics) in B1 containing 1% NGS. After washing in B1 buffer, the alkaline phosphatase activity was detected by using nitroblue tetrazolium chloride (337.5μg/ml) and 5-bromo-4-chloro-3-indolyl phosphate (175μg/ml) (Roche Diagnostics). Sections were mounted in Mowiol.

### Anatomical data acquisition and analysis

Images of PKCγ staining, BDA CST tracing, virus-labelled corticospinal innervation of cervical and lumbar spinal cord and retrograde tracing were acquired on Leica DMR microscope, under identical conditions of magnification, illumination and exposure. Gray level images were captured using MetaView software (Universal Imaging Corporation, Ropper Scientific, France). *In situ* hybridization slides were scanned with a Nanozoomer (Hamamatsu). Brightness and contrast were adjusted using Adobe Photoshop.

#### Laterality of the corticospinal, rubrospinal and reticulospinal trats based on retrograde tracing

FG sections were dehydrated with graded alcohol, immersed in xylene and mounted with Eukitt (Sigma). We first verified that FG was correctly injected on the right side of the cervical enlargement (C6) and that there was no leakage of the tracer accross the midline. Samples from 5 *Ntn1*^*lox/lox*^ and 6 *Shh::cre;Ntn1*^*lox/lox*^ mice fitted this criteriae (35% of the injected mice) and were included in the subsequent analysis. The structures of interest, i.e. medullary reticular formation, red nucleus and motor cortex, were captured on 3, 4 or 5 consecutive sections respectively with a Leica DMR microscope, under identical conditions of magnification, illumination and exposure. FG labelled neurons were segmented with ImageJ, using a selective threshold. After thresholding, we applied two algorithms of mathematical morphology: SKIZ (Skeleton by Influence Zones) and Watershed. For every animal, we reported the number of segmented neurons on each side (ipsilateral or contralateral to the injection site) of each structure (medullary reticular formation, red nucleus, motor cortex) and we calculated the ratio of ipsilateral segmented neurons over the total number of segmented neurons for each structure.

#### Quantification of labelled corticospinal axons in the white matter and distribution analysis of corticospinal synapses

were carried out from confocal images of spinal cord sections of 7 *Ntn1*^*lox/lox*^ and 7 *Shh::cre;Ntn1*^*lox/lox*^ mice injected unilaterally with AAV9-CAG-tdTomato virus in the motor cortex.

#### Quantification of labelled corticospinal axons in the white matter

Confocal laser scanning microscope (Leica SP8) equipped with a 63x objective (oil immersion, 1.4 Numerical Aperture) was used. Image stacks of 70 microns in width (144nm pixel size, 500 nm z-step) were acquired on controlled microscope stage and stitched into mosaic by the Leica LAS X software.

In control mice, corticospinal axons were detected mostly in the contralateral dorsal funiculus (1) and to a lesser extent in the ipsilateral dorsal funiculus (2) at C7 and L2. In mutant mice, corticospinal axons were detected in the contralateral dorsal funiculus (1), in the ipsilateral dorsal funiculus (2) in the ipsilateral lateral funiculus (3) and ipsilateral ventral funiculus (4) at C7 and in the first three first regions only at L2. Mosaics were therefore captured in the following regions of two consecutive sections: (1) and (2) at C7 and L2 for control animals and (1), (2), (3), (4) and (1), (2), (3) at C7 and L2 respectively for mutant animals. A region of interest was manually traced in each mosaic to define the area in which axons were segmented and counted. Segmentation was performed in three-dimensions (3D) by iterative thresholding and MESR algorithm^28,29^. User-defined constraints (i.e. minimum intensity threshold, minimum and maximum volumes) were included in the procedure. The segmentation was automated with macros calling functions from the ImageJ plugin 3DImageSuite.

#### Distribution of corticospinal synapses

Confocal laser scanning microscope (Leica SP5) equipped with a 63x objective (oil immersion, 1.4 Numerical Aperture) was used to acquire image stacks with pixel size of 160 nm and z-step of 300 nm. Image stacks of VGLUT1 were denoised with median filter.

At cervical and lumbar enlargement, the majority of corticospinal terminations were detected in the medial half of the laminae III-V (defined as the “dorsal” region) and the medial half of the laminae V-VII (defined as the “intermediate” region) in both *Ntn1*^*lox/lox*^ and *Shh::cre;Ntn1*^*lox/lox*^ mice. For each region of interest (ROI), three image stacks of 80 microns width and 8 microns depth were acquired on the ipsilateral and contralateral sides of the grey matter.

Segmentation in both channels was performed following 3D iterative procedure described above. Colocalization was detected in 3D by 3DImageSuite ImageJ plugin as overlap between segmented objects from the two channels.

For each section, we calculated the density of colocalization per ROI as the average of colocalization densities of the 3 sampling regions from this ROI. The densities of colocalization per ROI of the two consecutive sections were then averaged. The relative distribution of colocalization among the 4 ROIs (ispilateral dorsal, ipsilateral intermediate, contralateral dorsal, contralateral intermediate) was obtained by dividing the colocalization density of each ROI by the sum of the densities of the 4 ROIs. This allowed us to compare the relative distribution of the corticospinal synapses (VGLUT1/Tomato colocalization) among the 4 ROIs between animals independently from tracing differences.

### 3D analysis of the CST

2 *Ntn1*^*lox/lox*^ and 3 *Shh::cre;Ntn1*^*lox/lox*^ mice were injected unilaterally with AAV9-CAG-tdTomato virus in the motor cortex (cf. Anterograde tracing of the CST).

#### Tissue preparation

After perfusion, brains and spinal cords were collected and post-fixed in the same fixative overnight at 4 °C. Samples were pretreated with methanol before immunostaining. Samples were dehydrated with 50% (1h30), 80% (1h30) methanol/PBS and 100% methanol for 2×1h30. Samples were chilled at 4°C and bleached overnight in 6%H_2_O_2_ in methanol at 4°C. Samples were rinced in methanol, and rehydrated with 100% (2×1h30), 80% (1h30), 50% (1h30) methanol/PBS and PBS 1X (1h30).

#### Whole-Mount Immunostaining and Sample clearing

Samples were first incubated at room temperature on a rotating shaker in a solution (PBSGT) of PBS 1X containing 0.2% gelatin and 0.5% Triton X-100 (Sigma-Aldrich) for 3 days. Samples were next transferred to PBSGT containing 0.1% saponin and anti-DsRED (1/500, Living Colors, 632496) and placed at 37°C with rotation for 10 days. This was followed by six washes of 1h in PBSGT at room temperature. Next, samples were incubated in PBSGT containing 0.1% saponin and a donkey anti-rabbit conjugated to the Cy3 fluorophore (1/200, Jackson Immuno Research) for 2 days at 37°C with rotation. After six washes of 1h in PBSGT at room temperature, samples were cleared using a methanol clearing protocol, adapted from iDISCO+ protocol^30^ found at https://transparent-human-embryo.com.

#### 3D imaging and image processing

3D imaging was performed with an ultramicroscope using Inspector Pro software (LaVision BioTec). 3D volume and movies were generated using Imaris software. Crystals that might form from secondary antibody precipitation were eliminated using the surface tool by creating a mask around each volume.

### Electrophysiology procedures

The electrophysiology protocol was performed on 5 *Ntn1*^*lox/lox*^ and 9 *Shh::cre;Ntn1*^*lox/lox*^ adult mice from the group of animals that underwent the behavioral tests.

#### Intracortical stimulation of motor cortex

Anesthesia was induced with intraperitoneal injection of chloral hydrate (400mg/kg) to render the animal unresponsive to paw pinch. Animals were then placed in a stereotaxic frame. Body temperature was maintained at 37°C by a heating pad. A craniotomy was made over the hindlimb area of M1.

We used home-made bipolar tungsten electrodes (250 μm apart, impedance=6.5KΩ). Electrode penetration was made perpendicular to the pial surface at 1mm depth. The stimulation was applied in the hindlimb motor area (between 1 to 2.0mm lateral to bregma, between 0,5 to 1.5 caudal to bregma).

10 to 20 stimulations (40ms duration train of 0.6ms monophasic pulses at 350Hz) were delivered every 10 sec by a constant current stimulator (Digitimer, Ltd). For a given site, we started at low current and increased the amplitude until the lowest current that consistently produced a motor effect (in >50% of trials). A maximal current of 25mA was used; if no response was evoked at this intensity, the site was considered nonresponsive and the electrode was moved to another site (within the hindlimb motor area of the same hemisphere). The same procedure was then performed on the motor cortex of the opposite hemisphere.

#### Electromyography recordings

We recorded electromyographic (EMG) responses from the gastrocnemii muscles bilaterally evoked by the stimulation of one motor cortex. 2 homemade silver recording electrodes were inserted 5mm apart in left and right gastrocnemii muscles. The proper placement of the electrodes was controled by noting increased EMG activity upon passive plantar flexion. The EMG recordings were made with a differential AC amplifier and then digitized at 12.5 kHz and analysed with Spike2 program (CED). Raw EMG was smoothed and rectified after recording with Spike 2.

#### EMG Analysis

Data are presented as mean of EMG obtained from 10 to 20 stimulations of one or the two motor cortices (7 out of 14 mices). Proportion of each type of responses (“strictly” contralateral, “strictly” ispilateral and bilateral) to stimulation are calculated on effective motor contraction only.

### Training and behavioral testing

The behavioral studies were performed on 15 *Shh::cre;Ntn1*^*lox/lox*^ and 14 *Ntn1*^*lox/lox*^ control mice aged between 8 and 12 weeks; male and female mice were used. We verified that the repartition of male and female did not differ between groups (Chi2, p= 0.893). All the behavioral tests were performed blindly to the genotype. For the first three days, the mice were habituated to being handled by the experimenters in order to limit stress. Mice were then tested with a partial SHIRPA protocol (grasping, clasping and auditory tests were performed, and whisker state was evaluated) in order to rule out major neurological abnormalities.

#### The open field test

was used to evaluate spontaneous activity and locomotion: mice were placed in the center of a 0.25-m^2^ arena and allowed to explore freely for 5 minutes. During this time, they were tracked and recorded with a camera fixed above the arena, and the total walking distance was calculated with Topscan software.

#### The Ladder test

apparatus (Locotronic) consists of a 124 cm × 8 cm corridor with a floor composed of 78 bars each 1 cm apart. The mice were made to cross the corridor and the number of slips of the forelimbs, hindlimbs and tail was automatically detected by 158 infrared captors placed on the corridor walls (sampling frequency 1000 Hz). The test was repeated three times for each mouse and the sum of errors throughout the three trials was calculated. This test evaluates the precision and coordination of limb positioning.

#### Treadmill

Mice were placed on a transparent treadmill (14 cm × 6 cm) moving at 12 cm/s. After a short training session, the mice had to run for ten seconds, during which period the positioning of their paws was recorded by a camera fixed under the apparatus. The proportion of symmetric and asymmetric strides was calculated after excluding frames in which the mouse was not running.

#### Rotarod

The accelerating Rotarod (BIOSEB) consists of a horizontal rod 3 cm in diameter, turning on its longitudinal axis. The training phase consisted of walking on the rod at a rotational speed varying from 4 to 40 rpm for one minute. The mice were then subjected to four trials in which the speed of rotation increased gradually from 4 rpm to 40 rpm over 5 min. Time spent on the rod was recorded and averaged for the 4 trials. The test was repeated three days in a row with the same procedure, except that the training session was performed only on the first day.

#### Reaching exploratory behavior

When placed in a new environment, as a glass cylinder, mice engage in “reaching” exploratory behavior, in which they contact walls with their forepaws^18,31^. This contact can be made with the two paws simultaneously (symmetric movement) or independently (asymmetric). Ten reaching movements were recorded with a video camera and then examined frame-by-frame to calculate the number of asymmetric and symmetric movements.

#### Obstacle avoidance test

This test allows to evaluate adaptative locomotion^18,31^. It consists in placing the mouse on an obstructed treadmill, on which 2 obstacles were mounted one small (5mm height) and one large (10mm). The treadmill was moving at 10 cm/s for 2 minutes (acclimatization phase) and 15 cm/s for the 2 following minutes. The mice were accustomed to the task for two days and were recorded with a video camera on the third day. The number of alternate steps (asymmetric movements) and hops (symmetric movements) over the obstacles was calculated for both the forelimbs and the hindlimbs during the 15cm/s session. If the mouse was pausing before avoiding the obstacle, the movement was not considered in the analysis.

### Statistical analysis

Data were analyzed with SPSS statistical software version 24.0 (Chicago, Illinois, USA). Normality in the variable distributions was assessed by the Shapiro-Wilk test. Furthermore, the Levene test was performed to probe homogeneity of variances across groups. Variables that failed the Shapiro-Wilk or the Levene test were analyzed with nonparametric statistics using the one-way Kruskal–Wallis analysis of variance on ranks followed by Nemenyi test post hoc and Mann–Whitney rank sum tests for pair-wise multiple comparisons. Correlations were estimated with the Spearman method. Variables that passed the normality test were analyzed with ANOVA followed by the Bonferroni *post hoc* test for multiple comparisons, or with Student’s *t* test when comparing two groups. Paired data were analyzed by repeated-measures ANOVA with two factors, followed by the Bonferroni *post hoc* test for multiple comparisons. Correlations were estimated with the Pearson method. Categorical variables were compared using Pearson’s χ2 test or Fisher’s exact test.

## Supplementary Materials

**Movie S1**. 3D pyramidal decussation in a control mouse

**Movie S2**. Altered pyramidal decussation in a *Shh::cre;Ntn1*^*lox/lox*^ mouse (#1)

**Movie S3**. Altered pyramidal decussation in a *Shh::cre;Ntn1*^*lox/lox*^ mouse (#2)

**Movie S4**. *Ntn1*^*lox/lox*^ mouse performing alternate movements during treadmill locomotion

**Movie S5**. *Shh::cre;Ntn1*^*lox/lox*^ mouse performing alternate movements during treadmill locomotion

**Movie S6**. *Ntn1*^*lox/lox*^ mouse performing alternate movements during obstacle avoidance

**Movie S7**. *Shh::cre;Ntn1*^*lox/lox*^ mouse performing abnormal symmetric movements during obstacle avoidance

**Movie S8**. *Ntn1*^*lox/lox*^ mouse performing alternate movements during exploratory reaching

**Movie S9**. *Shh::cre;Ntn1*^*lox/lox*^ mouse performing abnormal symmetric movements during exploratory reaching

**Fig. S1**. Corticospinal tract guidance is disrupted at the pyramidal decussation in *Shh::cre;Ntn1*^*lox/lox*^ mice

**Fig. S2**. Expression of *Ntn1* is preserved in the forebrain of *Shh::cre;Ntn1*^*lox/lox*^ mice

**Fig. S3**. In the midbrain of *Shh::cre;Ntn1*^*lox/lox*^ mice, *Ntn1* depletion is not strictly limited to the floor plate

## General

We acknowledge France Lam for confocal imaging (IBPS, France), Morgane Belle (Institut de la Vision, France) for light-sheet imaging, Alexis Bemelmans (CEA, France) for producing the AAV9-CAG-tdTomato virus, Isabelle Caillé for reviewing the manuscript and PHENO-ICMice (ICM, France) Core supported by 2 “Investissements d’avenir” (ANR-10-IAIHU-06 and ANR-11-INBS-0011-NeurATRIS) and the “Fondation pour la Recherche Médicale”.

## Author contributions

O.P., A.C., E.R. and I.D. designed the study. O.P. and M.P.M. performed the behavioral analysis. O.P., Q.W. and M.P.M carried out the 2D anatomical analysis of the CST and O.P. performed the 3D anatomical analysis. N.H. implemented the segmentation procedure for confocal images. M.P.M. performed the genotyping. O.P. and F.M. performed the EMG recordings. N.S. carried out the FG injection. S.R.P. and J.A.M.B. performed the in situ experiments. M.D. carried out the segmentation procedure for retrograde labelling and performed the statistical analyses. O.P., E.R. and I.D. wrote the manuscript. Q.W., F.M., N.H., S.R.P, J.A.M.B, C.G., M.V., A.T., P.F., and AC substantively revised the manuscript.

## Supplementary Materials

**Movies (S1 to S9)**

### Movie S1. 3D pyramidal decussation in a control mouse

3D movie of a cleared brainstem of a *Ntn1*^*lox/lox*^ adult mouse in which the corticospinal tract originating from the left motor cortex is labeled with tdTomato. At the pyramidal decussation, all the labelled corticospinal axons turn dorsally and cross the midline to project into the dorsal funiculus of the contralateral spinal cord.

### Movie S2. Altered pyramidal decussation in a *Shh::cre;Ntn1*^*lox/lox*^ mouse (#1)

3D movie of a cleared brainstem of a *Shh::cre;Ntn1*^*lox/lox*^ adult mouse in which the corticospinal tract originating from the left motor cortex is labeled with tdTomato. At the pyramidal decussation, some of the labelled corticospinal axons turn dorsally and cross the midline, following the normal trajectory while the remaining axons do not cross the midline and project into the ipsilateral ventral spinal cord.

### Movie S3. Altered pyramidal decussation in a *Shh::cre;Ntn1*^*lox/lox*^ mouse (#2)

3D movie of a cleared brainstem of another *Shh::cre;Ntn1*^*lox/lox*^ adult mouse in which the corticospinal tract originating from the left motor cortex is labelled with tdTomato. In this mutant, some corticospinal axons turn dorsally, leaving the main tract just before the pyramidal decussation to project aberrantly into the dorsal funiculus of the ipsilateral spinal cord. Many axons remain ventral and project also into the ipsilateral spinal cord, while a few axons follow the normal trajectory and project into the dorsal funiculus of the contralateral spinal cord.

### Movie S4. *Ntn1*^*lox/lox*^ mouse performing alternate movements during treadmill locomotion

Extract of a video recorded during the treadmill locomotion task showing a *Ntn1*^*lox/lox*^ adult mouse. This mouse displays normal alternate movements with the forelimbs and hindlimbs when running on the treadmill. Video speed was reduced 6 times for better visualization.

### Movie S5. *Shh::cre;Ntn1*^*lox/lox*^ mouse performing alternate movements during treadmill locomotion

Extract of a video recorded during the treadmill locomotion task showing a *Shh::cre;Ntn1*^*lox/lox*^ adult mouse. This mouse displays normal alternate movements with the forelimbs and hindlimbs when running on the treadmill. No hopping can be detected. Video speed was reduced 6 times for better visualization.

### Movie S6. *Ntn1*^*lox/lox*^ mouse performing alternate movements during obstacle avoidance

Extract of a video recorded during the obstacle avoidance task showing a *Ntn1*^*lox/lox*^ adult mouse avoiding the high obstacle (1cm) with alternate movements of the forelimbs and the hindlimbs. Video speed was reduced 6 times for better visualization.

### Movie S7. *Shh::cre;Ntn1*^*lox/lox*^ mouse performing abnormal symmetric movements during obstacle avoidance

Extract of a video recorded during the obstacle avoidance task showing a *Shh::cre;Ntn1*^*lox/lox*^ adult mouse avoiding the high obstacle (1cm) with synchronous movements of the forelimbs and the hindlimbs. Video speed was reduced 6 times for better visualization.

### Movie S8. *Ntn1*^*lox/lox*^ mouse performing alternate movements during exploratory reaching

Extract of a video recorded during the exploratory reaching task showing a *Ntn1*^*lox/lox*^ adult mouse performing a typical exploratory behavior, contacting the walls with alternate movements of the forelimbs. Video speed was reduced 6 times for better visualization.

### Movie S9. *Shh::cre;Ntn1*^*lox/lox*^ mouse performing abnormal symmetric movements during exploratory reaching

Extract of a video recorded during the exploratory reaching task showing a *Shh::cre;Ntn1*^*lox/lox*^ adult mouse performing an abnormal symmetric exploratory movement with the forelimbs. Video speed was reduced 6 times for better visualization.

**Figures (S1 to S3)**

