## Supplementary Figure 1 for "Loss of floor plate Netrin-1 impairs midline crossing of corticospinal axons and leads to mirror movements"

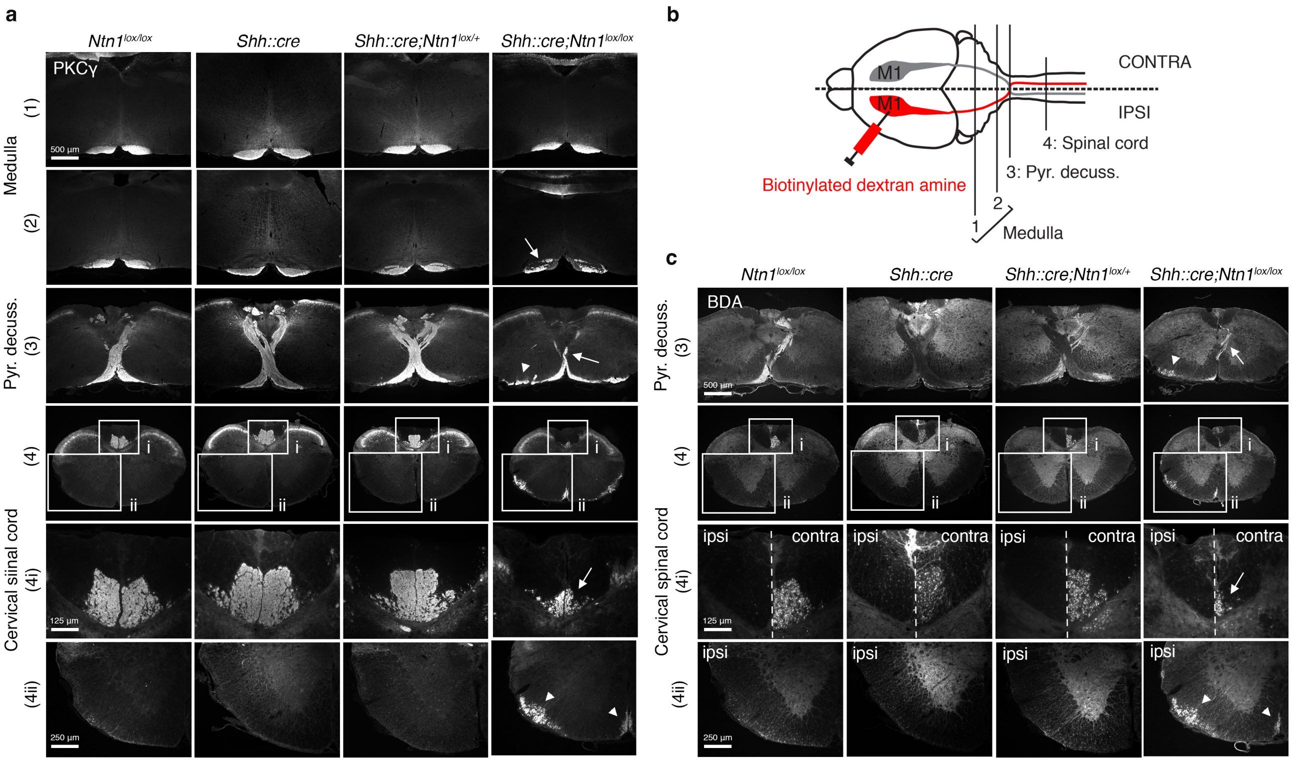


### Supplementary Figure 1: Corticospinal tract guidance is disrupted at the pyramidal decussation in *Shh::cre;Ntn1*$^{lox/lox}$ mice

*a*, Coronal sections of adult *Ntn1*$^{lox/lox}$ (n=4), *Shh::cre* (n=1), *Shh::cre;Ntn1*$^{lox/+}$(n=1) and *Shh::cre;Ntn1*$^{lox/lox}$ mice (n=5) stained with anti-PKCγ, a marker of the CST, at different levels: (1) upper and (2) lower medulla, (3) pyramidal decussation, and (4) cervical spinal cord. The CST is similar in all genetic background in (1). In (2), the CST is less fasciculated in *Shh::cre;Ntn1*$^{lox/lox}$ (arrow) compared to *Ntn1*$^{lox/lox}$, *Shh::cre*, or *Shh::cre;Ntn1*$^{lox/+}$ mice. In (3), pyramidal decussation of *Shh::cre;Ntn1*$^{lox/lox}$ is thinner (arrow) compared to *Ntn1*$^{lox/lox}$, *Shh::cre*, or *Shh::cre;Ntn1*$^{lox/+}$ mice and the CST spreads laterally (arrowhead). In (4), less axons are detected in the dorsal funiculi of *Shh::cre;Ntn1*$^{lox/lox}$ (4i, arrow, dotted line is the midline) and some axons are seen in ectopic positions in the ventromedial and lateral funiculi (4ii, arrowheads). **b**, Schematic showing the CNS levels for the coronal sections presented in a and c and strategy for unilateral BDA labelling of the CST. **c**, Same coronal sections as presented in a with revelation of the BDA tracer. (3): Fewer corticospinal axons cross the midline in *Shh::cre;Ntn1*$^{lox/lox}$ mice (arrow) compared to *Ntn1*$^{lox/lox}$, *Shh::cre*, or *Shh::cre;Ntn1*$^{lox/+}$ mice and the remaining axons are detected in the ipsilateral ventral white matter (arrowhead). In (4), less axons are detected in the contralateral dorsal funiculus in *Shh::cre;Ntn1*$^{lox/lox}$ mice (4i, arrow). Some ectopic CST axons are detected in the ipsilateral ventromedial and lateral funiculi (4ii, arrowheads).

BDA: biotinylated dextran amine. Pyr. decuss. = pyramidal decussation. ipsi, contra are respectively ipsilateral and contralateral sides with respect to the injection side.
