## Supplementary Figure 2 for "Loss of floor plate Netrin-1 impairs midline crossing of corticospinal axons and leads to mirror movements"

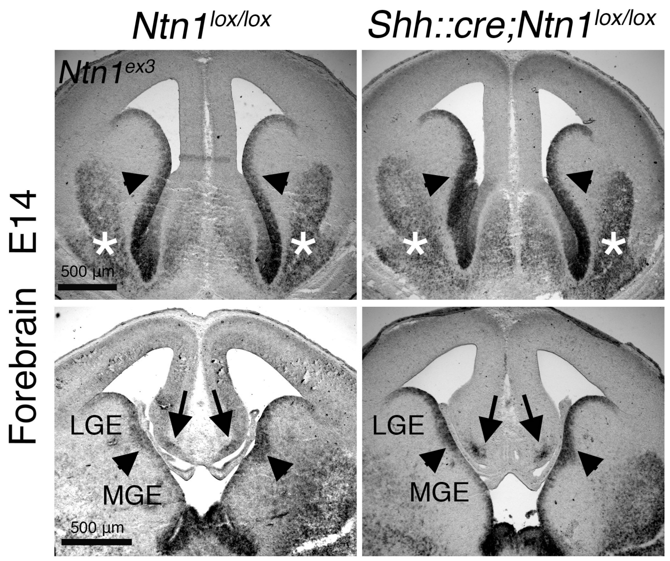


**Supplementary Figure 2: Expression of *Ntn1* is preserved in the forebrain of *Shh::cre;Ntn1$^{lox/lox}$* mice**

Coronal sections of E14 forebrain of *Ntn1$^{lox/lox}$* (n=3) and *Shh::cre;Ntn1$^{lox/lox}$* mice (n=3) (top: rostral, bottom: caudal level). At E14, Ntn1 mRNA is mostly expressed in the ventricular zone of the ganglionic eminences (arrowheads), in the ventral forebrain (white star) and in the glial wedges (arrows). This distribution is similar between *Ntn1$^{lox/lox}$* and *Shh::cre;Ntn1$^{lox/lox}$* mice. *Ntn1$^{ex3}$*: riboprobe against the exon 3 of the mouse *Ntn1* mRNA ; LGE/MGE: lateral/medial ganglionic eminence.
