## Supplementary Figure 3 for "Loss of floor plate Netrin-1 impairs midline crossing of corticospinal axons and leads to mirror movements"

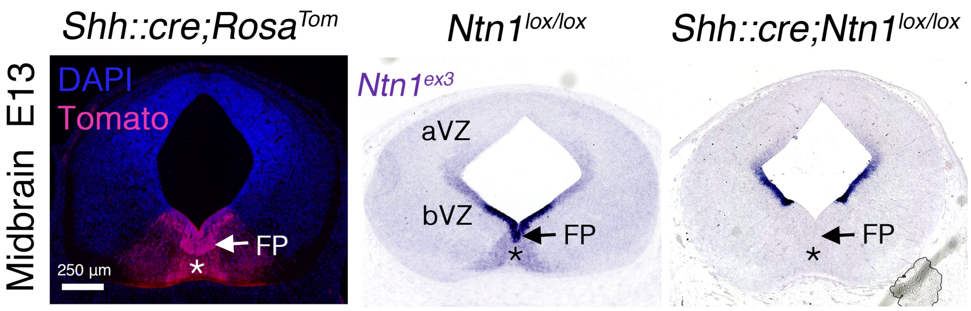


**Supplementary Figure 3: In the midbrain of *Shh::cre;Ntn1*$^{lox/lox}$ mice, *Ntn1* depletion is not strictly limited to the floor plate**

Coronal sections of E13 midbrain of *Shh::cre;Rosa*$^{Tom}$ mice (n=2), *Ntn1*$^{lox/lox}$ (n=3) and *Shh::cre;Ntn1*$^{lox/lox}$ mice (n=3). Left: In the midbrain, at E13, Tomato is expressed in the floor plate (FP, arrow), in the ventral part of the basal plate of the ventricular zone (bVZ, arrowhead), and in the midbrain dopaminergic neurons (star), which are *Shh* derivatives. Middle: *Ntn1* is highly expressed in the FP (arrow) and in the basal plate of the ventricular zone (bVZ). Lower expression is detected in the alar plate of the VZ (aVZ) and in midbrain dopaminergic neurons (star). Right: In *Shh::cre;Ntn1*$^{lox/lox}$ mice, compared to *Ntn1*$^{lox/lox}$ mice, *Ntn1* is depleted in the FP and the ventralmost part of the bVZ but its expression is intact in the more dorsal part of the bVZ and in the aVZ. Note that *Ntn1* is depleted in midbrain dopaminergic neurons (star). *Ntn1*$^{ex3}$: riboprobe against exon 3 of the mouse *Ntn1* mRNA; FP: floor plate; aVZ: alar plate ventricular zone; bVZ: basal plate ventricular zone.
